# A tale of two cores: a quantitative comparative analysis of the institutional and the open access models for flow research technology platforms

**DOI:** 10.1101/138552

**Authors:** A.M. Petrunkina, R. Schulte

## Quinquennium of ‘sorting for Cambridge‘

Biomedical specialist core facilities are major part of institutional infrastructure in academia and play important role in research by creating access to well established and emerging technologies and being a source of valuable expertise. In each particular case, the extent, range and content of services they provide are determined by the technology specialism, demands of the users and institutional/overarching policies.

In 2012, two flow cytometry core facilities embarked on a new journey in development of their services to research community in Cambridge. Both facilities started from the same initial level: mainly providing services in flow cytometry and cell sorting, each equipped with two sorters, staffed with two laboratory personnel and headed by a manager. While both cores provided the services in cell sorting, their approach to service provision, operations, access and funding model differed. In this brief case study we apply a mathematical formalism suggested by Petrunkina and Filby (2017) for utilization and efficiency metrics of sorting service in context of two different service provision models: institutional (IM) and centralised (CM) and discuss further avenues of facilities development.

The first important step both facilities made from the start was an attempt of a coordinated operation as interdependent parts of a joint service resource while remaining autonomous in terms of budget, policies and operational management. Rather than acting in competition for limited funding and resources, they applied for joint grants, operated coordinated tender exercises, implemented joint recruitment process, coordinated staff development, rotation for staff and backed up each other operations. This initial approach was highly successful. A core partnership received funding for capital equipment four times (initial investment, capacity expansion (sorters), capacity expansion (analysers) and new technology analysers). The contingency of operations was ensured by having identical equipment and staff trained in all technologies. However, the different access policy and different charging structure has defined further development and a divergence of initially closely coordinated operations into a casually associated collaboration.

During five years (2012-2017), both cores delivered high amount of cell separation services. A quinquennial review has identified a shortage in capacity for provision of sorting services in the institutional facility, despite their best efforts to sustain the existing level of services and adapt them to the persisting high demand. Due to change of landscape on campus (arrival and departure of new groups) it appears premature to initiate any new recruitment before the organisational changes are completed. How to balance the current shortage of staff capacity with expected fluctuations in demand? To find the best ways of dealing with it, one needs to conduct a quantitative comparative analysis of both models.

## Emerging divergence: origins

The institutional facility had a well organised set up, highly efficient on their existing scale of operations. It was funded by a major strategic award, and therefore was not exposed to high pressure of becoming sustainable in the short-term. All service contracts were funded by the major grant funding body, as was the salary of the lab manager, and half of each technician salary was funded by the central University funds. The facility has to recover only 1 FTE of the technical post (50% for each technician) and running costs/consumables. Lack of financial pressure, together with alignment to the overarching institutional strategy (such as compartmentalisation of focussed highly specialist services within one particular technology, e.g. imaging, cytometry, proteomics) has defined the direction of development. Moreover, the access was restricted to researchers connected in some way to their current location (tenants of their or nearby buildings, for example or their collaborators), thus the user base remained approximately constant due to usual turnover.

In developing the services they concentrated on enhancing capacity (ensuring provisions of sorting services, keeping abreast of the specialist technology development and adapting existing protocols and machines to particular researcher’s applications, such as index sorting, focussing on maximising the quantity of high quality services in this particular specialist area without expanding the portfolio of services (we will call that in depth growing of specialist operations). Over the period of 5 years, the institutional facility has provided services in a specialist flow cytometry range, including cell sorting, users training, analysis support, technical maintenance, advice and consultation, experimental support and primary data analysis. They have acquired new equipment, mainly to replace older kit and for capacity reasons (a sorter matching the specification of the first sorter, an analyser of largely same specification and one new analyser). In 2017, the institutional unit (IU) is staffed by three staff, has two sorters and seven analysers (plus associated equipment) and spread over three laboratories.

In contrast, the centralised facility has been immediately exposed to extremely high level of financial pressure. For years it has been operating at loss. It had no central funding or expectations to receive such, all service contract costs had to be met from Departmental reserves or from cost recovery. It has been equipped with outdated machines. In 2012 the resource has been awarded an external infrastructure funding sufficient for two staff posts. Given these conditions, its operational strategy has focussed on providing open access, expanding user base, increasing volume of operations, and enabling flexible fulfilment – agility and dynamic responsiveness to researcher’s requirements. Furthermore, it focussed on expanding portfolio of services, evolving new services and adding value to them while simultaneously increasing the capacity and facilitating specialist applications (we will call it in depth and breadth grow of specialist operations). Above and beyond to the services provided by the first facility, the centralised facility has evolved provision of new services over 5 years: imaging, high content and high throughput analysis/cellomics, magnetic enrichment/depletion, phenotyping, late operations, sample processing and added value/contract research work. These services can be offered at the access only, collaborative or comprehensive level, depending on the requirements of the researchers.

Such a wide-ranged approach allowed the second core to attract a large number of new users and collaborators, to utilise alternative ways of applying for funding (in addition to aforementioned joint applications, another four had successful outcome, and the initial infrastructure award has been increased subsequently). Enhancement of the infrastructure both in terms of staffing and machines, combined with the enrichment of service diversity enabled a qualitative transformation of a specialist core facility into a science technology/research platform (STP/RTP) by 2016. The centralised platform (STP) is staffed by ten staff, equipped with six sorters, nine analysers, kit for high content analysis, four specialised microscopes, magnetic separator and has integrated other specialised facilities such as tissue culture suite and general wet space, occupying 5 laboratories. In the next part of the study we will focus on the exclusive review of cell sorting services without reporting the output of other services.

## Shortage of capacity: perception or fact?

Figure 1 illustrates the output of sorting services in both institutional core facility and campus-wide platform given in accountable hours billed to the grants. It is immediately apparent from this figure that capacity in both services appears to be saturated over last two years. But can one claim that the saturation of output signals the shortage of capacity or is that something which can be improved by increasing efficiency and utilizing the existing resources better? Can one deduce from the reports that the central platform is having a nearly double capacity as the institutional unit? Or can one imply they are actually performing below capacity because their infrastructure is 300% of the institutional facility but they deliver ‘only’ twice as much sorting?

**Figure 1.**
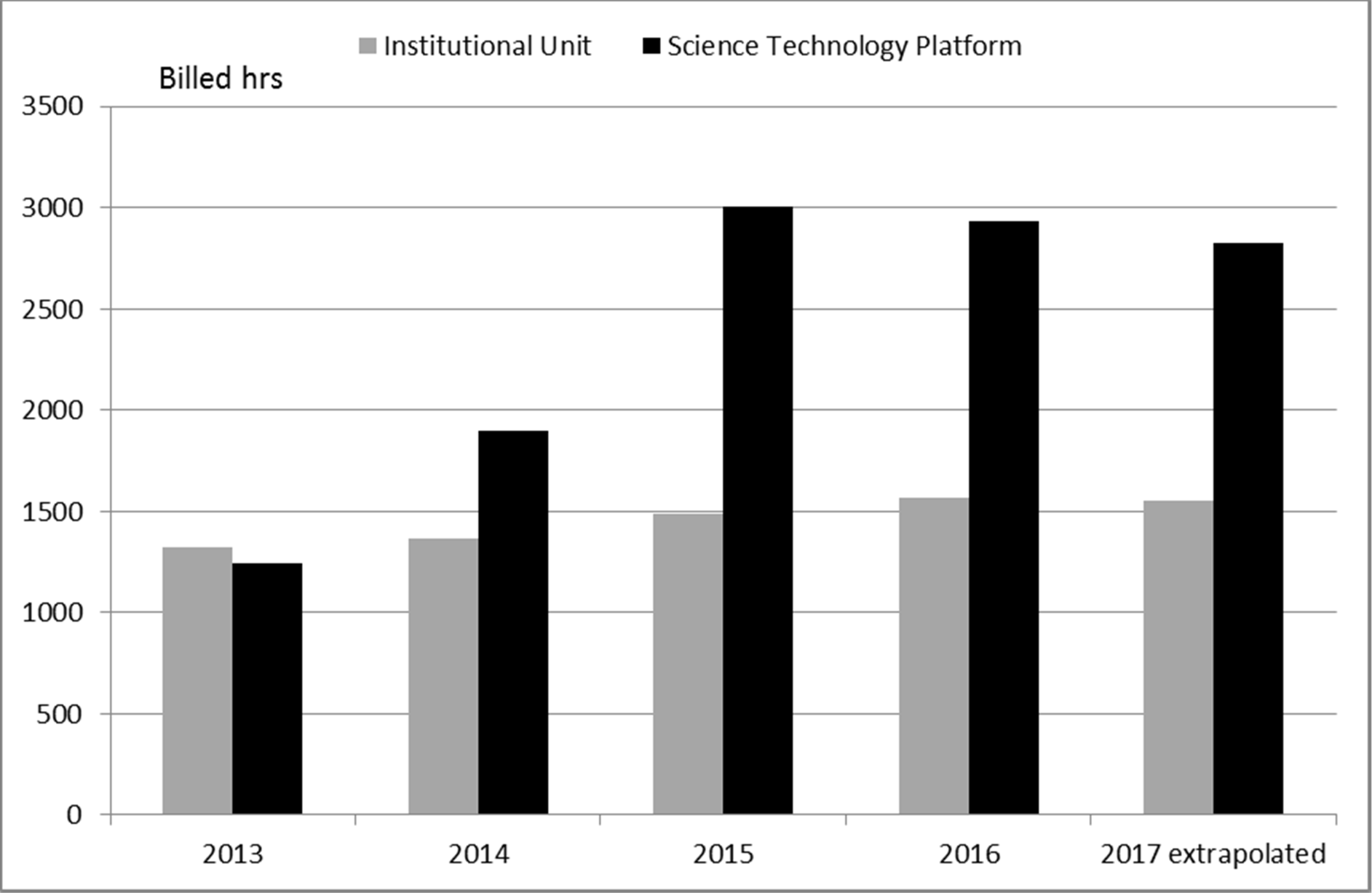
Performance of sorting service (output in billable hours of sorting)

The answer to any of these questions cannot be given without further analysis. The fact is that it is impossible to compare the absolute outcomes of service outside the context of operational strategy, organisational policy and market analysis. Thus, to conduct such an analysis, we suggest using the metrics of capacity and utilization set out in the formalism published by Petrunkina (2013).

Using this algorithm, one will be proceeding stepwise. First step is to express the nominal staff numbers and their time dedicated to the sorting in fulltime equivalents. This information required to conduct these calculations can be extracted from job descriptions, the length of employment over year in question and whether they work(ed) full time or part-time. The resulting data (Figure 2) strongly indicate that amount of staffing in the institutional facility and their allotted sorting time remained constant over the years (3 FTE staffing, 1.5 FTE sorting time, slightly fluctuating during short-staffing periods in 2013 and 2016 when facility was understaffed for 1.5 and 2 months respectively. Both nominal staffing and the allotted sorting time increased in the central platform (8.18 FTE and 3.14 FTE, respectively). These data clearly demonstrate that a linear approach to describing capacity (nominal staffing/hours of work) is erroneous and would have obscured the fact that the staff-based sorting capacity of the STP is only about the double of that of the IU due to distribution of various duties across the service portfolio and their ‘sorting portion’ (Figure 3).

**Figure 2.**
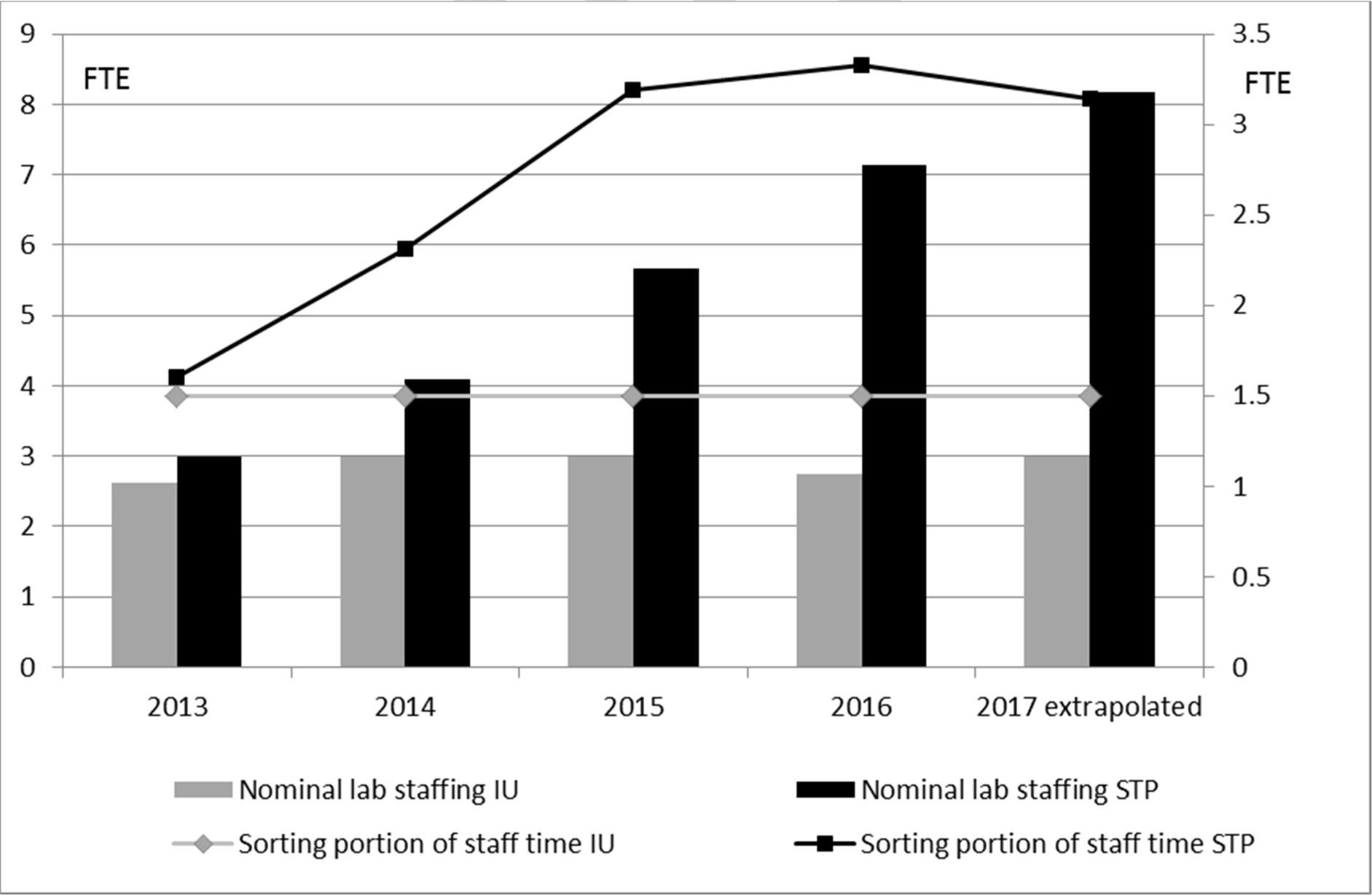
Staffing in the laboratories, expressed in nominal and sorting full time equivalents (FTE).

**Figure 3.**
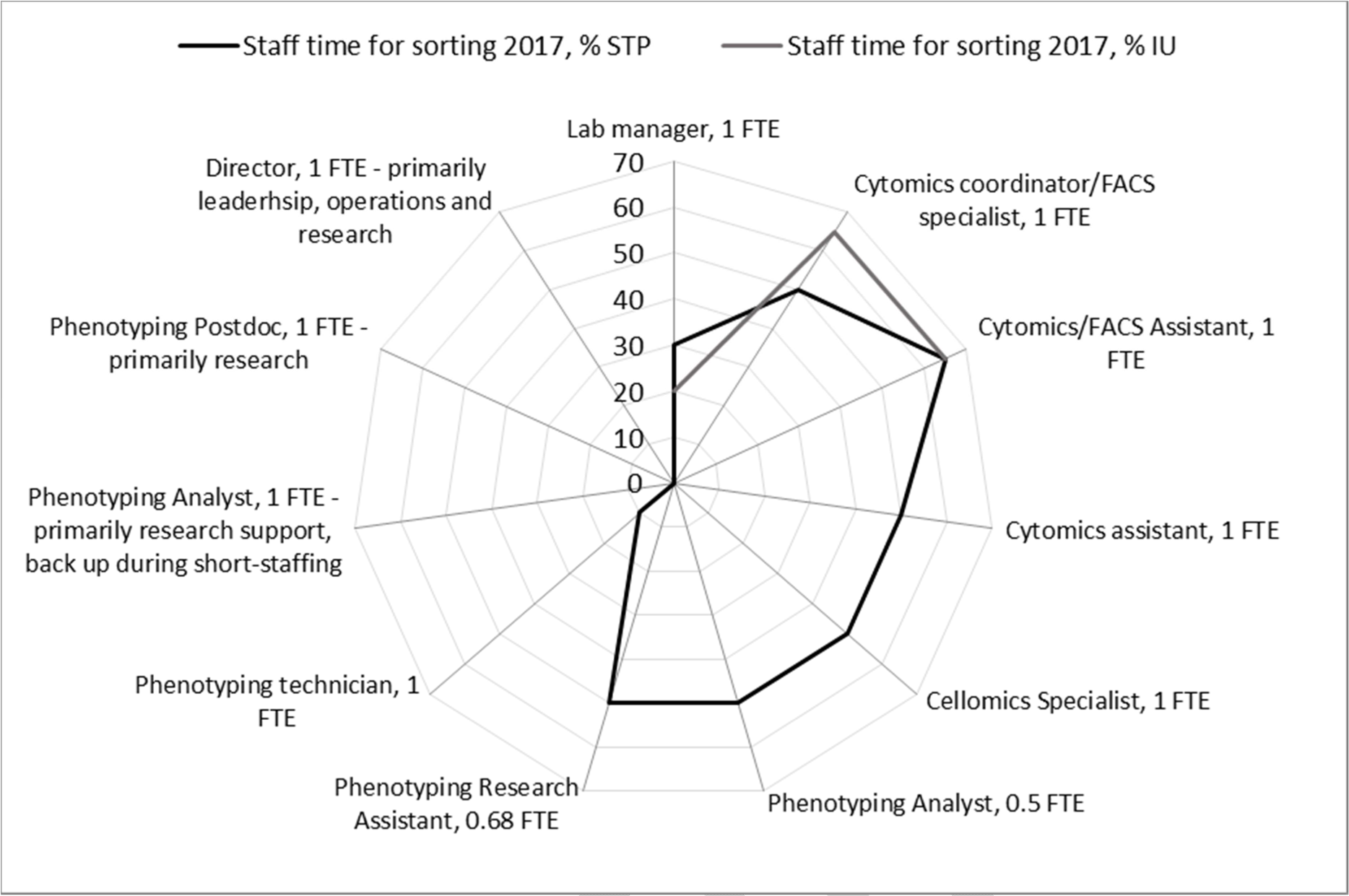
Sorting component of different roles in Institutional Unit and Research/ Science Technology Platform.

However, the reported billable sorting hours (1566 vs 2937 in 2016) are still not directly comparable, even after FTE-adjustment. Both facilities report output data as billable and not bookable time, thus it is not possible to link these data directly to the time physically available for operation that will include non-billable operative time (preparation, set up, decontamination, calibration etc). As an additional internal discrepancy, the institutional facility provides data as pure sorting time without set up and controls runs (time range 15-45 minutes for each run depending on sort complexity), and that set up time needs to be added up to the billable/deducted from the available booking time. The centralised platform provides the data with the set up because it is billable time according to their policies but they have higher incidence of cancellations due to clinical reasons (working on human samples), therefore they intentionally book more time than they expect to bill which affects the perceived capacity. Both these aspects need to be incorporated. Beyond that, the output in billable hours does not tell us whether the output (comparable or not) has exhausted the capacity in each case.

The next step therefore would be to calculate the total booking time available for operation of sorters. It will be equal to the product of cumulative sorting portion of staff time in FTE units multiplied with their annual working time in hours, less preparation and shutdown time to relate exclusively to the time available for billable sorting service delivery. Then, these figures have to be adjusted for efficiency. Efficiency of utilization will correct for the differences between number of working days and number of actual days worked in the lab, which is less by any authorised absences like holidays, sickness, course attendance etc and breakdowns/refurbishments (Petrunkina, 2013). In this particular case, for the sake of simplicity we do not conduct a detailed retrospective analysis of efficiency but assume that both units operate at optimal efficiency of working time level (80%, Petrunkina and Filby, 2017). That brings us to the 1627 hrs capacity of sorting in total for IU and 3125 hrs for STP). The final step in our calculations would be to adjust these bookable hours to expected billable hours (Figure 4).

**Figure 4.**
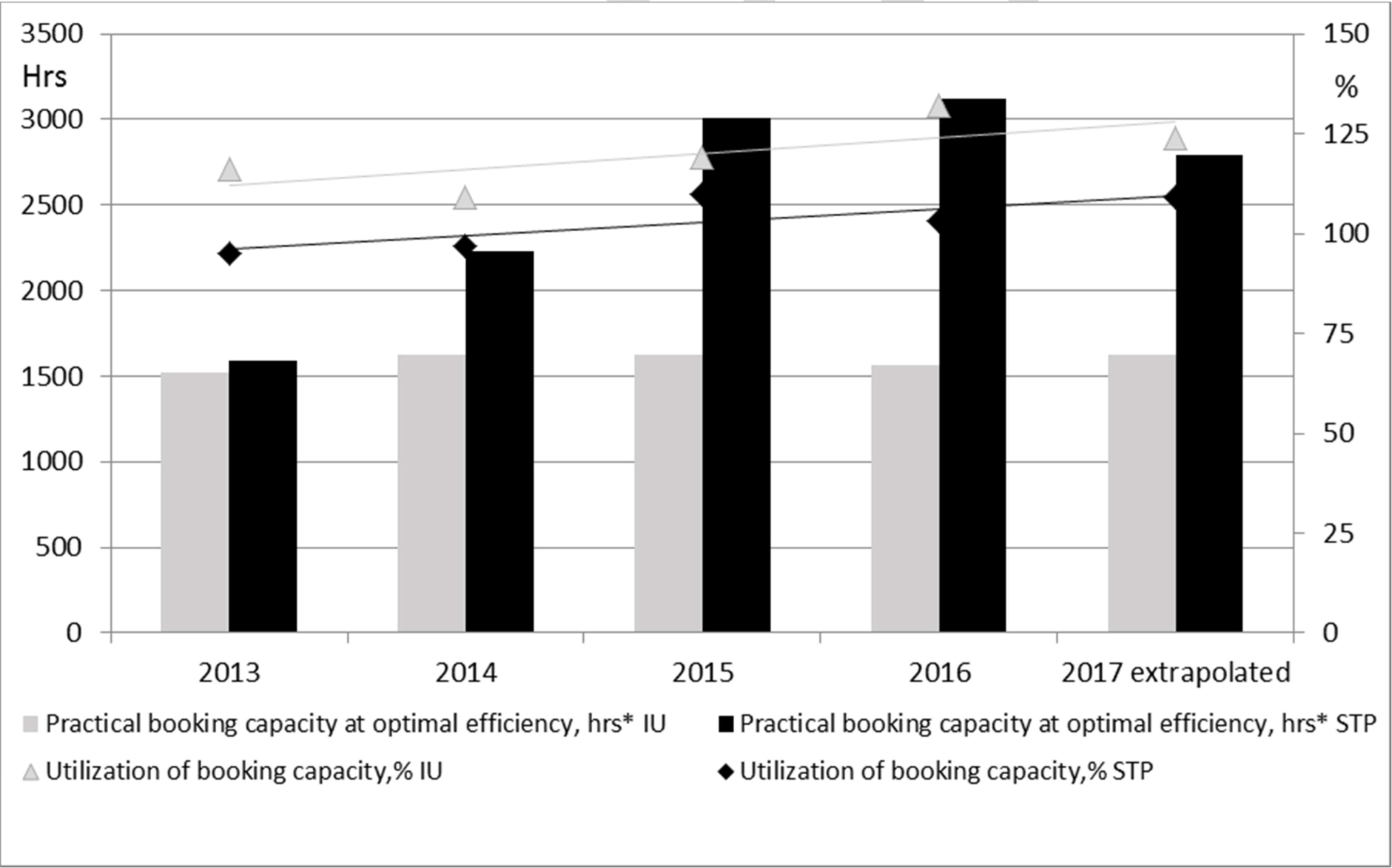
Standardised practical booking capacity at optimal efficiency in IU and STP and utilization of efficiency over the 5 years period.

## Current and projected capacity benchmarking

The derived practical billable capacity will be 1250 hours for institutional facility under consideration of set-up times. For central platform, under consideration of clinically-caused cancellations, the practical billable capacity will be ^~^2800 billable hours (Figure 4). Once arrived at that stage, we can further deduce that IU utilized ^~^132% of their capacity in billable sorts, and that STP utilized ^~^104% of their capacity in billable sorts in 2016. The extrapolation for 2017 based on initial reports suggests that IU and STP, respectively, are likely to utilize 125% and 110% of available capacity.

What does it mean in plain English? It means that both facilities outperform in terms of utilization of available resources at optimal efficiency (s.f. Petrunkina and Filby, 2017). It also shows that the dynamic of demand on sorting evolved in a similar way in both units (^~^3-5% persistent annual increase in capacity utilization, Fig. 4). Furthermore, it shows that the institutional facility is in a ‘near-crisis’ state with respect to the heavy and persisting demand on its core staff. Delivering at that level might mean that people may skip their lunches, neglect own professional development for the lack of time, not take their holidays and work long hours. While such dedication is highly commendable, helps to attain desired results in the short-term, it may also mean that the concerned staff will start looking for a less stressful job soon.

Central platform, while performing above the benchmark, is in an overall more beneficial position in terms of contingency for effective and efficient service. First, they ‘only’ run at 10% above the capacity. The tolerance threshold may stretch to this level, especially if the stress is not lasting permanently and a relief is expected. Due to the operational strategy of STP, a developed infrastructure and diversity of job descriptions, there is also higher level of adaptability and flexibility for contingencies: a specialist from different service area may be ‘conscripted’ into sorting duties which increases the sorting capacity range for period of time and absorb overtime.

## Conclusions

The quantitative analysis of the cell sorting service delivery for two facilities in Cambridge shows that the algorithm for benchmarking and standardised metrics can be fully applied to real situations. It does provide unequivocal evidence that both resources are run efficiently and performing above capacity with respect to provision of sorting services. It indicates, in addition, that the current capacity of the institutional facility is severely limited.

At the operational level, this case report is also expected to initiate discussions whether one should use the universally adjustable billable capacity metrics (according to the algorithm laid out in Petrunkina and Filby, 2017) for calculating and setting a uniform user access fee across sites/facilities? Should one use the adjusted booking capacity metrics for better coordinating/managing booking schedule and relieving the pressure on booking system, e.g. by allocating quota for ‘external users’ or acquiring new automated equipment?

At the strategic level, our results encourage a more general discussion to analyse critically the advantages and disadvantages of institutional (departmental) and centralised models.

Core facilities are key support infrastructures that provide specific technologies and expertise in cutting-edge technologies in an affordable manner. It has been suggested that the core facility concept ideally should be taken beyond single institutions towards institutional alliances (Meder et al., 2016). Such an approach would equip scientists with the major sophisticated and agile toolkit for the interdisciplinary approaches. In our day and age, it will soon become impossible and unaffordable to maintain the state of the art technologies at the institutional level, both in context of science and sustainability. At the same time, a local access to the state of the art services such as provided by institutional facilities is crucial.

Raising awareness of benefits associated with moving towards larger versatile or coordinated platforms/networks beyond single institutions (such as superior flexibility, agility, diversity, job satisfaction, compatibility of fully flexible service with work-life balance and ultimate staff retention) while acknowledging the need for conservation of advantages related to existing local access facilities and/or establishing new local outlets must be carefully balanced factors in such a discussion. A hybrid model combining best features of either model could be a future solution representing a first step towards alliances of institutional platforms.

